# Pupillometry signatures of sustained attention and working memory

**DOI:** 10.1101/2021.01.18.426969

**Authors:** Paul A. Keene, Megan T. deBettencourt, Edward Awh, Edward K. Vogel

## Abstract

There exists an intricate relationship between attention and working memory. Recent work has further established that attention and working memory fluctuate synchronously, by tightly interleaving sustained attention and working memory tasks. This work has raised many open questions about physiological signatures underlying these behavioral fluctuations. Across two experiments, we explore pupil dynamics using real-time triggering in conjunction with an interleaved sustained attention and working memory task. In Experiment 1, we use behavioral real-time triggering and replicate recent findings from our lab (deBettencourt et al., 2019) that sustained attention fluctuates concurrently with the number of items maintained in working memory. Furthermore, highly attentive moments, detected via behavior, also exhibited larger pupil sizes. In Experiment 2, we develop a novel real-time pupil triggering technique to track pupil size fluctuations in real time and trigger working memory probes. We reveal that this pupil triggering procedure elicits differences in sustained attention, as indexed by response time. These experiments reflect methodological advances in real-time triggering and further characterize an important biomarker of sustained attention.

## Introduction

Attention and working memory are closely linked. In particular, recent work has interleaved sustained attention and working memory tasks to demonstrate that attention and working memory fluctuate synchronously (deBettencourt et al., 2019). That is, lapses of sustained attention covary with lapses of working memory (Adam & deBettencourt, 2019). Various biomarkers of attention and working memory have been discovered, including pupil size, which may provide access and insight into the interrelationship between these mechanisms (Robison & Unsworth, 2018; Unsworth & Robison, 2016). In particular, these biomarkers may provide important insight into the synchronous cognitive dynamics of attention and working memory.

Extensive research has implicated pupil size as a key measure of attention, arousal and general task engagement (Clewett et al., 2020; Eldar et al., 2013; Gilzenrat et al., 2010; Joshi et al., 2016; Kahneman & Beatty, 1966). In particular, differences in pupil size have been implicated in sustained attention (Decker et al., 2020; Unsworth & Robison, 2016; van den Brink et al., 2016) and working memory tasks (Robison & Brewer, 2020; Robison & Unsworth, 2018; Unsworth & Robison, 2015; Zokaei et al., 2019). Thus, we adapted our procedure to continuously record pupil size while participants performed our interleaved sustained attention and working memory task.

In particular, we leveraged real-time triggering to examine the relationship between sustained attention and working memory, as well as whether pupil size can provide a continuous index of these cognitive processes. Real-time triggering is a procedure that automatically and adaptively designs experiments, contingent to fluctuations of cognitive state. We have previously employed this experimental approach by examining behavioral data in real time and monitoring for lapse-prone attentional states. We successfully detected moments when attention was especially high or low and inserted memory probes contingent to these attention fluctuations (deBettencourt et al., 2018, 2019). Other related work has developed similar triggering approaches derived from biomarkers instead of behavior (Chew et al., 2019; Hinds et al., 2013; Yoo et al., 2012). Real-time triggering can thus be a powerful approach for exploring the relationship between endogenous cognitive and physiological fluctuations.

The goal of this experiment is to explore how pupil size relates to sustained attention and working memory. In Experiment 1, we used real-time triggering derived from behavioral fluctuations of sustained attention. Behavioral real-time triggering tracked trial-to-trial fluctuations of response time and triggered working memory probes whenever prepotent responses were especially fast (i.e., lapsing attentional states) or slow (i.e., attentive states). In Experiment 2, we developed a novel real-time triggering procedure derived from fluctuations of pupil size. Pupil real-time triggering tracked trial-to-trial fluctuations of pupil size and triggered working memory probes whenever pupil sizes were especially large or small. These distinct and complementary real-time triggering procedures allow us to directly target extremely attentive and inattentive states, as measured by behavior and pupillometry.

### Experiment 1

The goal of Experiment 1 was to examine physiological signatures that underlie sustained attention and working memory, using an interleaved task.

## Methods

### Participants

Thirty-five people participated in Experiment 1 for $25 payment (20 female, mean 25.5 years). Two participants left the study early without completing it. One participant accidentally participated twice; their second session was excluded from analysis. Two participants were excluded for working memory performance that was worse than chance-level guessing. This resulted in a final sample size of *n*=30 participants. This sample size exceeds our target of twenty-four, based on prior work (deBettencourt et al., 2019). All participants reported normal or corrected-to-normal color vision and provided informed consent to a protocol approved by the University of Chicago Social & Behavioral Sciences Institutional Review Board.

### Apparatus

Participants were seated 75 cm away from an LCD monitor (120-Hz refresh rate). Stimuli were presented using Python and Psychopy.

### Stimuli

Stimuli were shapes, either circles (diameter = 1.5°) or squares (1.5°x1.5°). Each display consisted of 6 shapes at 4° eccentricity. The shape positions were consistent for all trials to minimize inter-trial visual transients. The color of each shape was one of nine distinct colors (red, blue, green, yellow, magenta, cyan, white, black, orange), and each display contained six shapes of random, unique colors. A central black fixation dot (0.1°) appeared at the center and turned white after a key press. For whole report working memory probes, a multicolored square (1.5°x1.5°) comprised of all nine colors appeared at each of the 6 locations, and the mouse cursor appeared at the central fixation position.

### Procedure

Participants completed an interleaved sustained attention and working memory task (Figure **1a**), adapted from a recent publication (deBettencourt et al., 2019). Critically, the sustained attention and working memory tasks relied on orthogonal dimensions of the stimuli: shape was the relevant dimension for the sustained attention task and color was the relevant dimension for the working memory task. The key difference from the previously published work was that the retention interval duration was increased (from 1 s to 2 s) to better capture slow pupillary responses. In total, participants completed 5 blocks of 800 trials. Due to time constraints, three participants completed only 4 blocks.

**Figure 1.**
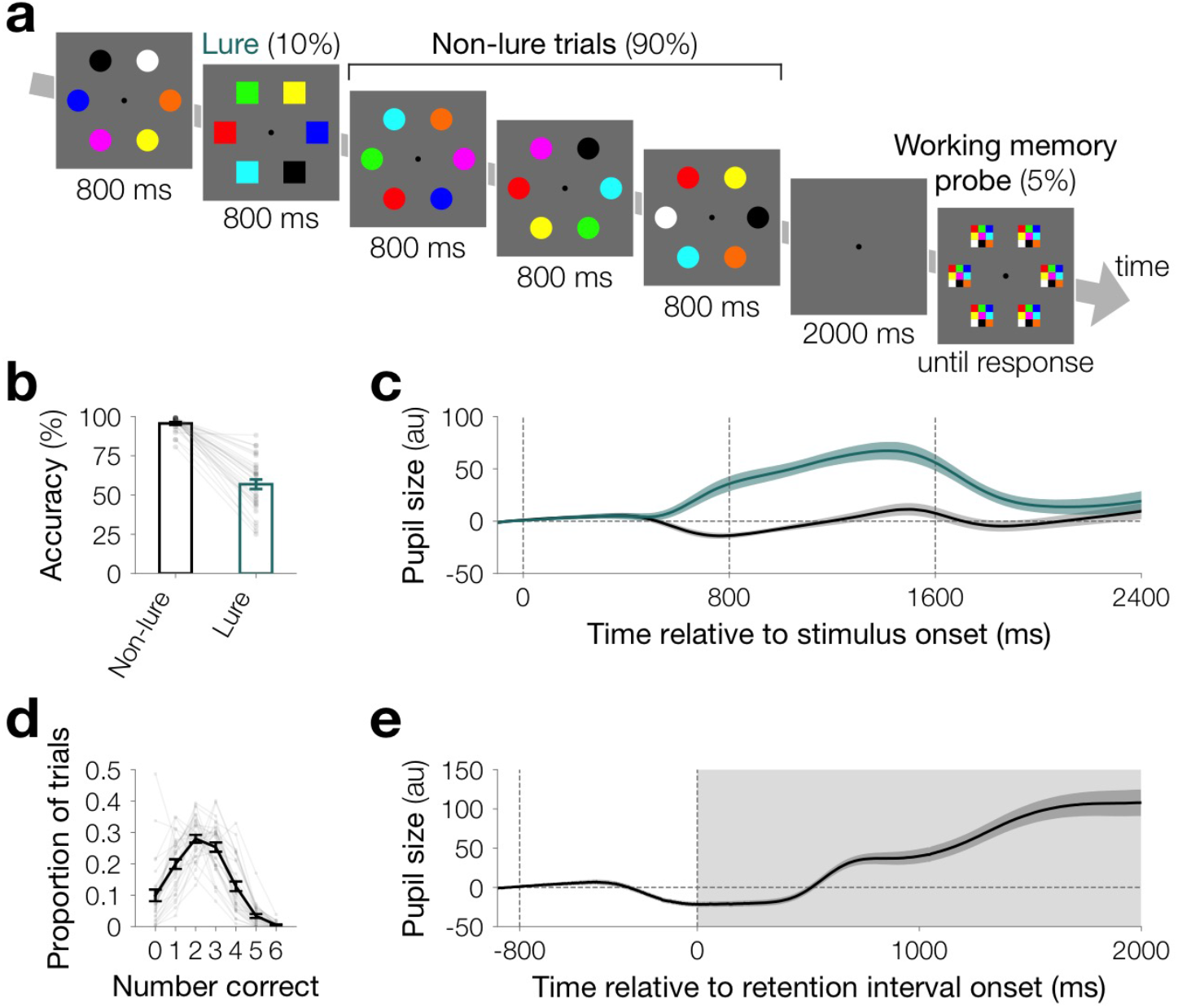
**a** Task design. Each trial was an array of six items (either circles or squares) of different colors. For the sustained attention task, participants responded to the shape, and for the working memory task participants reported the color. To encourage habitual responding, one of the shapes was much more frequent (10% lure trials, squares). For the whole report working memory task, after a 2 s delay participants selected the color of each item. **b** Sustained attention accuracy. Accuracy was higher for lure vs. non-lure trials (*p*<0.001), a key signature of sustained attention tasks. The height of the bars depict the average accuracy, and error bars are the standard error of the mean. Data from each participant are overlaid as small gray dots connected with lines. **c** Pupil size evoked by the sustained attention task. Lure trials (teal) evoked large pupil sizes than non-lure trials (black). Pupil size is plotted in arbitrary units (a.u.) relative to a 100 ms baseline prior to the stimulus onset. Each line is the population average timecouse, and the shaded area is the standard error of the mean. **d** Working memory performance. In the whole report color working memory task, performance on each trial ranged from 0 (no items correct) to 6 (all items correct). The black line depicts the average proportion of trials, and error bars are within-subject standard error of the mean. Data from each participant are overlaid as small gray dots connected with gray lines. **e** Pupil size during the working memory retention. Pupil size increased during the blank retention interval. Pupil size is plotted in arbitrary units (a.u.) relative to a 100 ms baseline prior to the stimulus onset. The black line is the population average timecourse, and the shaded area is the standard error of the mean.

In the sustained attention task, participants viewed displays consisting of 6 circles or 6 squares. Their task was to respond according to the shape of the stimuli: If the shapes were squares, they pressed the “s” key, and if the shapes were circles, they pressed the “d” key. To effectively manipulate sustained attention, one shape was especially prevalent (approximately 90% circles, 10% squares). To reduce stimulus transients, the stimuli were at fixed positions and remained on the screen for 800 ms with no inter-stimulus interval in all experiments.

In the working memory task, participants were asked to remember the colors of each of the 6 shapes that most recently appeared. Whole report color working memory probes appeared infrequently; approximately 4-5% of trials were probed. Memory probes were inserted randomly during the first 80 trials. For the rest of the block, memory probe trials were inserted contingent to response behavior afterwards (see real-time triggering procedure below). During a working memory probe, all shapes disappeared and the screen went blank and gray with only the central black fixation dot for a brief delay interval (2 s). Then, multicolored squares, which included all nine possible colors, appeared at each location. The participants selected one of the nine colors at each location using the mouse cursor before the screen would advance. After making a response at a particular location, a large black square appeared around the outside of that entire multicolored square. After the last response, the screen went blank again (1 s) before resuming the sustained attention task.

### Real-time triggering procedure

The goal was to use occasional working memory probes to link fluctuations in sustained attentional state with working memory performance. Rather than randomly inserting working memory probes, probes were “triggered” based on behavioral fluctuations of sustained attention. We tracked trial-by-trial fluctuations of sustained attentional state by continuously monitoring RT. For each trial *i*, we calculated a measure of attentional state using the average response time of the three most recent trials (*i*-3, *i*-2, *i*-1). We triggered memory probes whenever this measure of sustained attentional state, pre-trial RTs, exceeded fast or slow thresholds. In real time, we developed individually tailored and adaptively updated thresholds based on the cumulative mean and standard deviation (trials 1, 2,…, i-1). When pre-trial RTs exceeded one standard deviation away from the cumulative mean RT, we triggered a memory probe. Specifically, faster responses (faster than the mean RT minus the standard deviation) indicated a lapsing sustained attentional state, while slower responses (slower than the mean RT plus the standard deviation) indicated an attentive state. All working memory probes belonged to the frequent category (i.e., circles).

While the RT-based thresholds were the primary criteria for triggering, we had additional criteria to optimize the targeting of attentional state, based on behavioral responses and eyetracking measures. We initiated the real-time triggering procedure after the first 80 trials. To ensure a reliable pre-trial RT measure of attentional state, we required that participants had made responses to each of the three most recent trials. We also ensured that no probe trials had appeared in the three most recent trials. Finally, any eye blinks during the retention interval aborted the probe trial. Participants were not informed about the real-time triggering procedure.

### Eye tracking

We monitored pupil size and gaze position using a desk-mounted infrared eye tracking system (EyeLink 1000 Plus, SR Research, Ontario, Canada). Eye tracking data were binocularly sampled at 1000Hz, and head position was stabilized with a chin rest and calibrated with a 5-point calibration procedure. We report eye-tracking results for the left eye, but both eyes were highly reliably correlated (*r*=0.98). Pupil size data are expressed in arbitrary units, corresponding to the size of the pupil as measured by the infrared camera. Using gaze position, distance, and pixel size, we calculated the degree of visual angle from central fixation. We detected missing data, blinks and saccades (>0.5°) using an automatic artifact pipeline, developed in our laboratory for analyzing eye-tracking data collected during EEG experiments. One possible concern is that pupil size differences are driven by stimulus-specific differences in luminance of the categorical colors. However, these stimuli were presented peripherally at 4° eccentricity, and 6 of 9 colors appeared on every trial.

### Analysis

Behavioral performance was analyzed for the interleaved sustained attention and working memory task. Sustained attention performance to each trial was characterized using accuracy and RT. Accuracy to infrequent and frequent trials were combined into a single nonparametric measure of sensitivity of A’ and compared vs. chance (0.5). Whole report working memory performance was characterized as the number of items per trial for which the participants selected the correct color. We further modeled working memory abilities for each individual using a computational model previously developed by our laboratory (Adam et al., 2015; Hakim et al., 2020), in which working memory performance is described by attentional control (*α*), maximum working memory capacity (*K*_max_) and guessing. To examine the influence of our real-time triggering design, we calculated the trailing window RT (what served to trigger memory probes). We also calculated the working memory performance for each memory probe.

Pupil size was also analyzed for the sustained attention and working memory task. We calculated the average pupil size evoked by each stimulus, using a 100 ms pre-stimulus baseline. We compared the average pupil size elicited by lure and non-lure trials after artifact rejection. We also calculated the average pupil size evoked during the working memory retention interval after artifact rejection, using a 100 ms pre-stimulus baseline. To examine the pupil size elicited by real-time triggering, we calculated the pupil size of the stimulus that served as a memory probe. We examined the pupil size for all trials and also after artifact rejection.

### Statistics

Because some of the data violated the assumption of normality, all statistics were computed using a nonparametric random-effects approach in which participants were resampled with replacement 100,000 times. Null hypothesis testing was performed by calculating the proportion of the iterations in which the bootstrapped mean was in the opposite direction. Exact *p*-values are reported; *p*-values that were smaller than one in one thousand are approximated as p<0.001. Hypotheses were directional and thus one-sided unless otherwise noted. The mean and standard error of the mean are reported as descriptive statistics. Correlations were computed using the non-parametric Spearman rank-order correlation function. All data and code will be made available upon publication.

## Results

We first examined whether the interleaved sustained attention and working memory task with eye tracking (Figure **1a**) elicited established behavioral and pupillary signatures. Sustained attention accuracy was lower for lure trials compared to non-lure trials (acc_infreq_=56.86±3.04%; acc_freq_=95.59±0.85%; *p*<0.001; Figure **1b**), a well-characterized signature of sustained attention task designs. Lure trials evoked a larger pupillary response (*s*_non-lure_=−0.03±3.42, *s*_lure_=29.36±4.60; *t*=0–2400 ms; *p*<0.001; Figure **1c**). Even after controlling for accuracy, lure trials evoked a larger pupillary response (*s*_non-lure_=−0.51±3.43, *s*_lure_=26.20±4.97; t=0–2400 ms; *p*<0.001). Overall, sustained attention performance was well above chance (*A′*=0.86±0.01, chance=0.5, *p*<0.001). Working memory performance was calculated as the number of items correctly reported in working memory probes (*m*=2.24±0.11). We further characterized working memory performance distributions (Figure **1d**) using a computational model that has been previously developed by our laboratory. This computational model describes whole report working memory performance for each participant using two factors, attentional control (*α*=1.81±0.23) and maximum working memory capacity (*K*_max_=2.83±0.18). The blank retention interval evoked a large pupil response (*s*_ret_=49.55±8.12; t=0–2000 ms; Figure **1e**). Working memory performance (*m*) was positively correlated with pupil size during the retention interval (*s*_ret_) across individuals, such that higher accuracy was observed when the pupil was larger (*r*=0.62; *p*<0.001). These findings show that participants successfully performed the interleaved sustained attention and working memory task, and the task elicited established pupillary signatures.

Prior work has established that RT in similar sustained attention tasks tracks fluctuations of sustained attention. Specifically, faster responses in this task reflect worse attentional states, as when participants are responding more quickly, they are more likely to lapse (i.e., incorrectly respond to lure trials). We calculated the average preceding RTs over a trailing window of the 3 most recent trials. Indeed, we replicated prior work that faster responses precede lapses (*RT*_lapse_=324±40, *RT*_non-lapse_=444±48, *p*<0.001). That is, in this task, faster responses to non-lure trials index worse attentional states.

We used real-time triggering to deploy working memory probes at specific moments, contingent to fluctuations of attentional state. As participants performed the sustained attention task, we tracked attentional state in real time by continually monitoring fluctuations of RT. Then, we triggered working memory probes (Figure **2a**) during moments when the participant was especially fast (operationalized as inattentive moments) or slow (operationalized as inattentive moments). We adapted this real-time triggering procedure for concurrent eye tracking by extending the retention interval (2 s from 1 s) to accommodate the slower pupillary response and discontinuing any working memory probes if we detected a blink during the retention interval.

**Figure 2.**
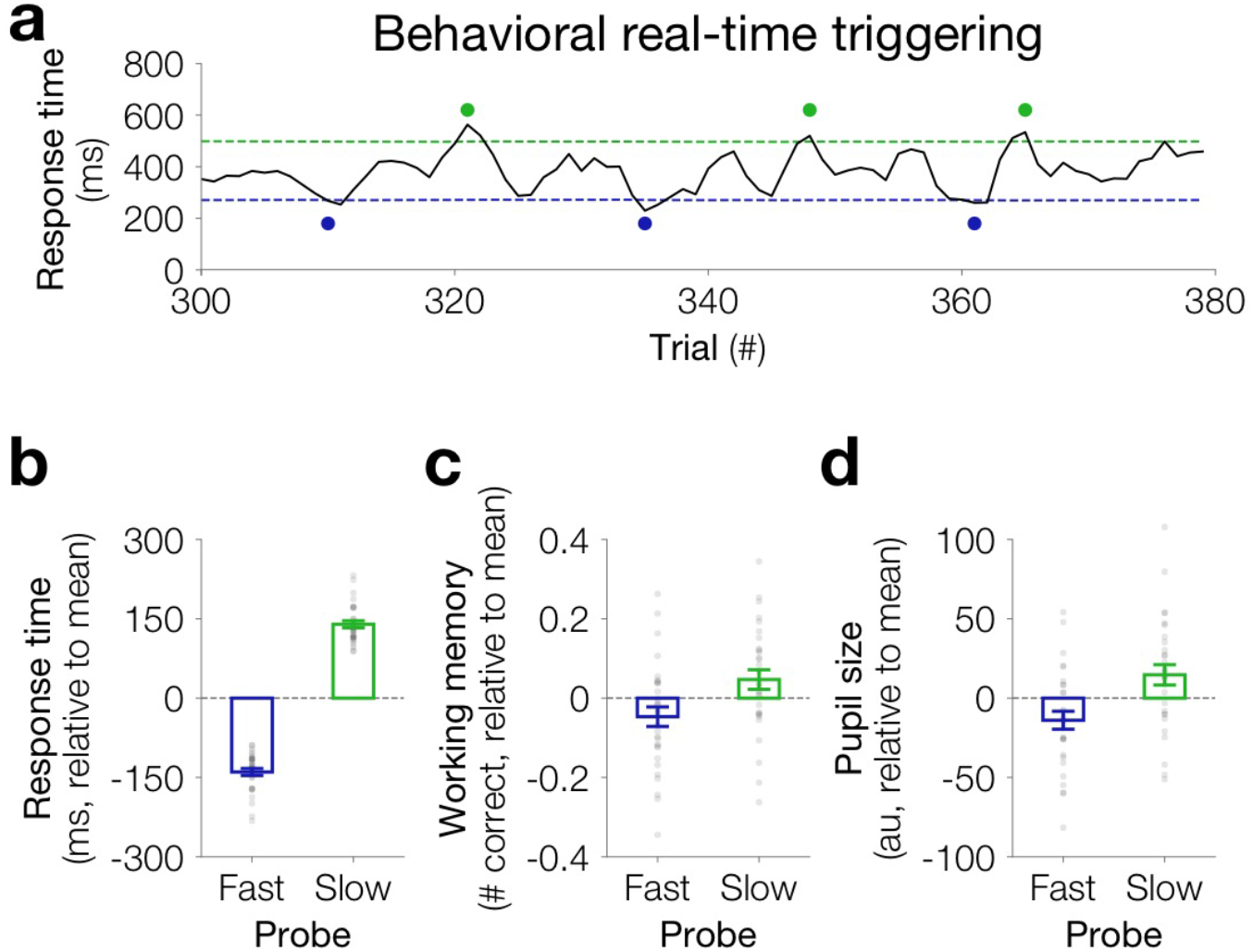
**a** Behavioral triggering. Behavioral fluctuations of RT (black line) were monitored in real time. We computed average RTs over a trailing window of the three most recent trials. Working memory probes (blue or green dots) were triggered whenever RT was faster than the fast threshold (blue, one standard deviation below average RT) or slower than the slow threshold (green, one standard deviation above average RT). **b** The real-time triggering procedure successfully detected faster RTs (blue) and slower RTs (green). The dashed gray line indicates the average RT before memory probes. **c** Working memory lapses concurrently with sustained attention. Participants remembered fewer items when working memory probes were triggered due to faster RTs (inattentive moments) versus slower RTs (attentive moments. The dashed gray line indicates the average working memory performance. **d** Pupil size differences covary with sustained attention. Average pupil size was computed over same interval as the trailing window for RTs. Pupil sizes were smaller when responses were faster (inattentive moments) versus slower (attentive moments). Bar heights depict the population average, and error bars are the standard error of the mean. Data from each participant are overlaid as small gray dots.

First, we investigated whether this real-time triggering manipulation was successful. Indeed, we successfully triggered trials when participants were responding fast or slow (RT_fast_=240±8 ms, RT_slow_=519±13 ms; Figure **2b**). That is, we continuously monitored behavior so as to detect attentive or inattentive moments.

Next, we examined whether attention fluctuated concurrently with working memory. If so, then participants would remember fewer items during inattentive moments. Indeed, we found that working memory performance was worse for fast-vs. slow-triggered trials (*m*_fast_=2.20±0.10, *m*_slow_=2.29±0.11; *p*=0.03; Figure **2c**). This behavioral pattern replicates the findings from our previous study, that sustained attention and working memory lapse concurrently.

Finally, we examined whether real-time triggering also captured differences in pupil size. Prior to fast-triggered working memory probes, pupil sizes were reliably smaller (f_fast_–9.12±5.22, *s*_slow_=9.05±5.45; *p*=0.04; Figure **2d**). This result was remained reliable even after excluding any trials with eye artifacts (*s*_fast_=-13.96±5.67; *s*_slow_=14.67±6.41; *p*=0.008). Thus, real-time triggering based on behavior revealed smaller pupil size during inattentive trials. Note that we are not claiming that RT and pupil size are redundant or directly equivalent measures of attentional state. Rather, that these moments detected by our real-time triggering method also detected moments with reliable differences in pupil size.

## Discussion

This experiment replicated the behavioral finding that sustained attention and working memory lapse concurrently, initially demonstrated in a prior publication from our laboratory (deBettencourt et al., 2019). We also extended these results to demonstrate that moments of low attention, operationalized as response time and detected by our automatic real-time triggering procedure, also reflected moments when the pupil size is smaller. These results also raise an important question about whether pupil size could serve as an independent and continuous index to track fluctuations of sustained attentional and/or working memory.

### Experiment 2

The goals of Experiment 2 were to develop a real-time triggering procedure based on moment-to-moment fluctuations of pupil size to explore the consequences of this triggering procedure for behavior.

## Methods

### Participants

Twenty-seven people participated in Experiment 2 for $25 payment (14 female, mean 23.9 years). Two participants left the study early without completing it. One subject was excluded due to multiple issues with data collection during the first two blocks of the task. This sample size matches our target of twenty-four, based on prior work (deBettencourt et al., 2019). All participants reported normal or corrected-to-normal color vision and provided informed consent to a protocol approved by the University of Chicago Social & Behavioral Sciences Institutional Review Board.

### Apparatus & Stimuli

Same as Experiment 1.

### Procedure

Same as Experiment 1. The difference was that the working memory probes appeared contingent to fluctuations of pupil diameter, not RT. In total, participants completed 5 blocks of 800 trials. For one participant, the response keys for the sustained attention task were swapped. For another participant, only data from the last four blocks was included in the analysis due to multiple errors in the first block.

### Real-time triggering procedure

Rather than randomly distributing working memory probes, memory probes were “triggered” based on real-time pupil size fluctuations. For each trial *i*, we calculated the average pupil size of the left pupil using a trailing window over the three most recent trials (*i*-3, *i*-2, *i*-1). We triggered memory probes whenever pupil sizes were especially small or large. In real time, we developed individually tailored and adaptively updated thresholds based on the cumulative mean and standard deviation of pupil size (from trials 1, 2,…, *i*-1). When the trailing window of pupil size exceeded one standard deviation away from the cumulative mean pupil size, a working memory probe was triggered. We initiated the real-time triggering procedure after the first 80 trials. We also ensured that no probe trials had appeared in the three most recent trials. In addition, any eye blinks during the retention interval aborted the probe trial. Participants were not informed that their pupils controlled when memory probes would appear.

### Eye tracking

For the first two subjects, eye tracking data were binocularly sampled at 1000Hz. To accelerate real-time computation, we switched to monocular tracking of the left eye for all subsequent subjects. All other details were the same as Experiment 1.

### Analysis

In addition to the analyses from Experiment 1, we examined the continuous relationship between sustained attention behavior and working memory behavior. To do this, we examined all working memory probes that were triggered for each participant, regardless of whether they were triggered due to large or small pupil sizes. For each participant, we correlated pre-trial RTs with the number correct on the working memory probe. To conduct statistics across participants, we *z-*transformed the *r*-values.

### Statistics

Same as Experiment 1.

## Results

Participants performed the interleaved sustained attention and working memory task with concurrent eye tracking. Sustained attention accuracy was lower for lure trials compared to non-lure trials (acc_infreq_=51.32±3.72%; acc_freq_=94.60±0.84%;*p*<0.001; Figure **3a**). Lure trials evoked a larger pupillary response (*s*_non-lure_=−2.29±3.48, *s*_lure_=27.68±6.48; *t*=0–2400 ms; *p*<0.001; Figure **3b**). Even after excluding trials with incorrect responses, lure trials evoked a larger pupillary response (*s*_non-lure_=−2.99±3.52, *s*_lure_=27.68±6.48; t=0–2400 ms; *p*<0.001). Overall, sustained attention performance was well above chance (*A′*=0.84±0.02, chance=0.5, *p*<0.001). Working memory performance was calculated as the number of items correctly reported in working memory probes (*m* = 2.07±0.13; Figure **3c**). We further characterized working memory performance distributions using a computational model with two factors, attentional control (*α*=1.31±0.17) and maximum working memory capacity (*K*_max_=2.75±0.14). The blank retention interval evoked a large pupil response (*s*_ret_=58.47±9.81; t=0–2000 ms; Figure **3d**). Working memory performance (*m*) was positively correlated with pupil size during the retention interval (*s*ret) across individuals (*r*=0.50; *p*=0.01). These findings replicated those from Experiment 1 and demonstrated that new pupil triggering task elicited established pupillary signatures.

**Figure 3.**
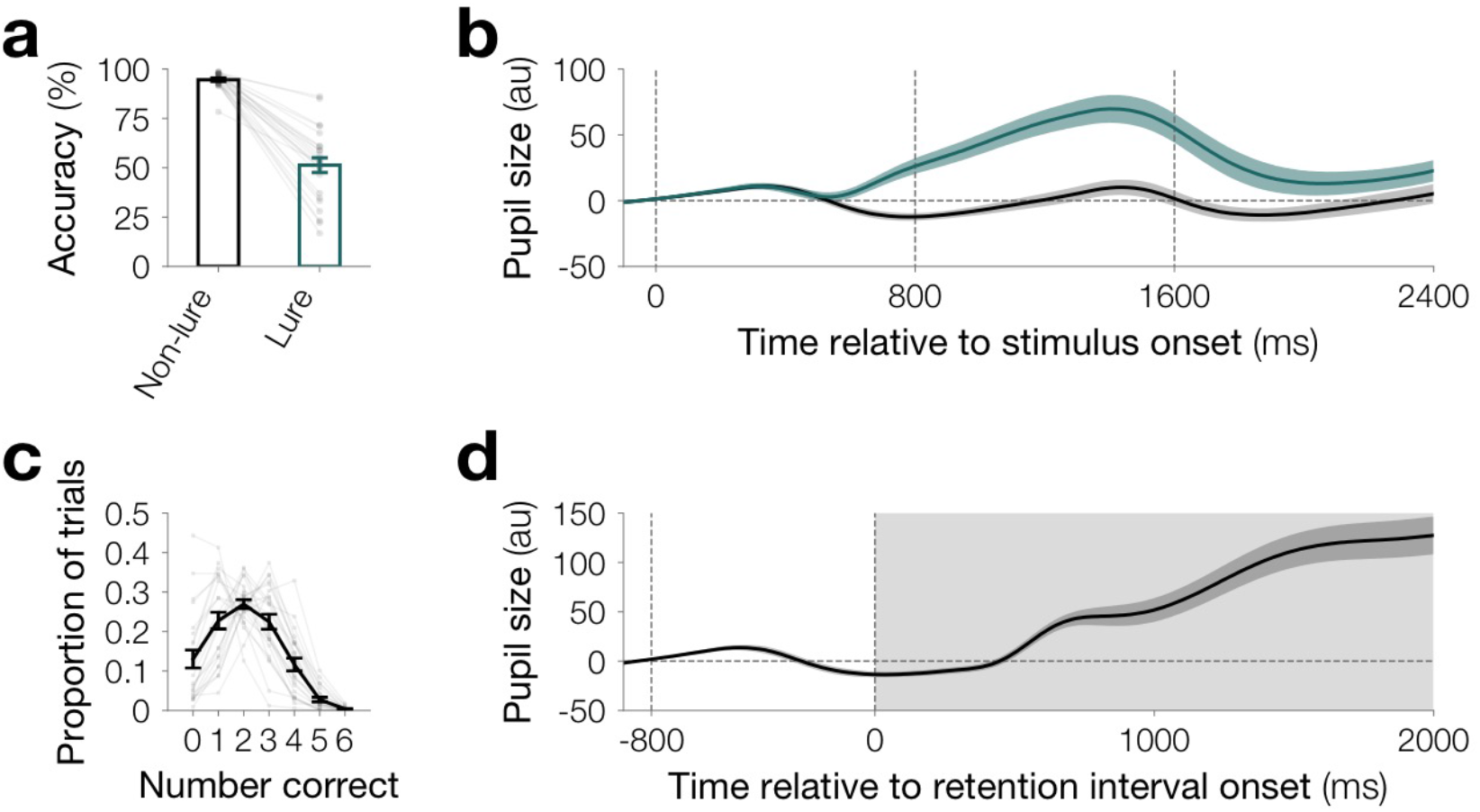
**a** Sustained attention accuracy. Accuracy was higher for lure vs. non-lure trials (*p*<0.001), a key signature of sustained attention tasks. The height of the bars depict the average accuracy, and error bars are the standard error of the mean. Data from each participant are overlaid as small gray dots connected with lines. **b** Pupil size evoked by the sustained attention task. Lure trials (teal) evoked large pupil sizes than non-lure trials (black). Pupil size is plotted in arbitrary units (a.u.) relative to a 100 ms baseline prior to the stimulus onset. Each line is the population average timecouse, and the shaded area is the standard error of the mean. **c** Working memory performance. In the whole report color working memory task, performance on each trial ranged from 0 (no items correct) to 6 (all items correct). The black line depicts the average proportion of trials, and error bars are within-subject standard error of the mean. Data from each participant are overlaid as small gray dots connected with gray lines. **d** Pupil size during the working memory retention. Pupil size increased during the blank retention interval. Pupil size is plotted in arbitrary units (a.u.) relative to a 100 ms baseline prior to the stimulus onset. The black line is the population average timecourse, and the shaded area is the standard error of the mean.

We also replicated the finding that RT is a signature of sustained attentional state. We calculated a measure of attentional state by averaging RT over a trailing window of the 3 most recent trials. Indeed, faster responses preceded lapses (*RT*_lapse_=311±38, *RT*_non-lapse_=372±42, *p*<0.001). Therefore, in this task, we again verified that faster responses reflected moments of worse sustained attention.

The critical difference is that Experiment 2 used pupil real-time triggering to deploy working memory probes at specific moments, contingent to fluctuations of pupil size (rather than RT). As participants performed the sustained attention task, we continually monitored fluctuations of pupil size via real-time eye tracking. Then, we triggered working memory probes (Figure **4a**) during moments when the pupil sizes were especially small or large.

**Figure 4.**
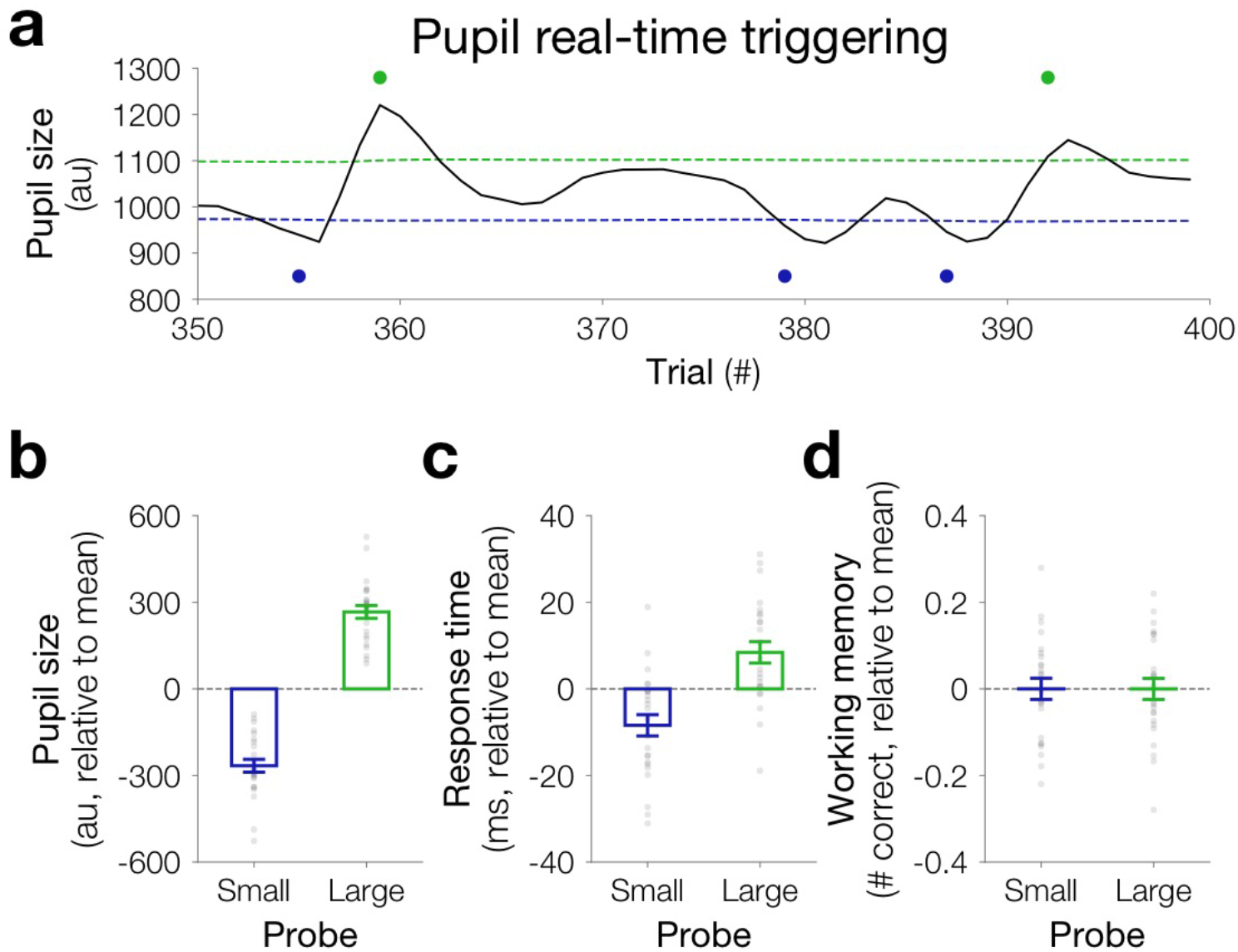
**a** Pupil real-time triggering example. Pupil size fluctuations (black line) were monitored in real time. We computed averaged pupil size over a trailing window of the three most recent trials. Working memory probes (blue or green dots) were triggered whenever pupil size was smaller than the small threshold (blue, one standard deviation below average pupil size) or larger than the large threshold (green, one standard deviation above average pupil size). **b** Real-time triggering successfully detected extreme differences in pre-trial pupil sizes. The average pupil size was computed over the preceding three trials before memory probes. As expected by the real-time triggering design, the procedure successfully detected moments of small (blue) and large (green) pupils. The dashed line indicates the average pupil size before memory probes. **c** Sustained attention covaries with pupil size differences. RT was measured over the three trials preceding each memory probe. Small (blue) vs. large (green) pupils covaried with sustained attention behavior (RTs, averaged over the three preceding trials). Smaller pupils covaried with worse sustained attention (i.e., faster RTs). The dashed line indicates the mean pre-trial RTs. **d** Pupil size differences do not covary with working memory fluctuations. Working memory was operationalized as the number correct on the whole report working memory probe. Small (blue) and large (green) pupils did not influence working memory performance. The dashed line indicates the mean working memory accuracy for real-time triggered memory probes. The height of the bar depicts the population average, and error bars are within-subject standard error of the mean. Data from each participant are overlaid as small gray dots.

First, we investigated whether this real-time triggering manipulation was successful. Indeed, we successfully triggered trials when participants’ pupils were large or small (*S*_small_=−194±21, *s*_large_=243±24; Figure **4b**), relative to mean pupil size. That is, we successfully continuously monitored for deviations in pupil size as participants performed the sustained attention task.

We next examined whether our measure of sustained attention, RT, fluctuated concurrently with pupil size. If so, then participants would be responding more quickly when pupil size was smaller. Indeed, we found that RTs were faster for small-vs. large-triggered trials (RT_small_=336±10 ms; RT_large_=353±11 ms; *p*<0.001 Figure **4c**). This demonstrates the complement of the relationship from Experiment 1, that real-time triggering based on pupil size can elicit reliable differences in RT.

We then examined whether pupil triggering would also predict differences in working memory performance. We did not observe a reliable effect of pupil triggering on working memory behavior (*m*_small_==2.08±0.13; *m*_large_=2.08±0.14 ms;*p*=0.50; Figure **4c**).

Finally, we examined whether these results were consistent with Experiment 1, by considering the relationship between attention and working memory behavior. For each participant, we examined all triggered working memory probes, regardless of whether they were triggered due to small or large pupils. For each participant, we correlated our behavioral measure of sustained attention (i.e., pre-trial RTs) with our behavioral measure of working memory (number correct for that memory probe). Across participants, there was a modest but reliably positive relationship between pre-trial RTs and working memory performance (mean *r*=0.06±0.03, *p*=0.03). Thus, these results confirmed our prior demonstrations of synchronous fluctuations between sustained attention and working memory behavior.

## Discussion

In Experiment 2, we developed real-time pupil triggering procedure to track fluctuations of a physiological measure as participants performed an interleaved sustained attention and working memory task. Pupil real-time triggering was sufficient to elicit but reliable changes in sustained attention behavior but did not give rise to differences in working memory accuracy. This suggests ways in which real-time triggering can be used to covertly and continuously track cognitive states and to disentangle attention and working memory.

### General Discussion

Across two experiments we measured pupil size as participants performed an interleaved sustained attention and working memory task. We triggered working memory probes in real time at precise moments, either based on RTs to the sustained attention task (Experiment 1) or pupil size (Experiment 2). In both experiments, we observed that the sustained attention task and the retention interval elicited canonical pupil responses. In the first experiment, we replicated the general relationship between sustained attention and working memory, showing that WM performance was lower when subjects were performing worse on the sustained attention task. In the second experiment, pupil real-time triggering elicited differences in sustained attention, but not working memory. That is, via complementary automated real-time triggering systems both experiments linked tonic fluctuations in pupil size with fluctuations of sustained attention.

#### Real-time triggering

This study demonstrates the potential for real-time triggering techniques from multiple behavioral and/or physiological indices within the context of the same task. Through behavioral real-time triggering, (Figure **5a**), we can investigate the consequences of extreme differences in attentional state (operationalized by speed of responding) and elicit reliable differences in pupil size (Figure **5b**) and working memory. Alternatively, pupil real-time triggering can be used to obtain a sensitive assay of how variation in pupil size track variations in cognitive performance by emphasizing the trials with the biggest differences in pupil size (Figure **5c**) and elicit modest yet reliable differences in responding (Figure **5d**). This work opens the potential for designing experiments where real-time triggering is derived jointly from behavior and physiological measures, either during moments of convergence (e.g., slow responses and large pupils) or divergence (e.g., slow responses and small pupils). By comparing the behavioral consequences of multiple indices, we can begin to disentangle complex cognitive dynamics.

**Figure 5.**
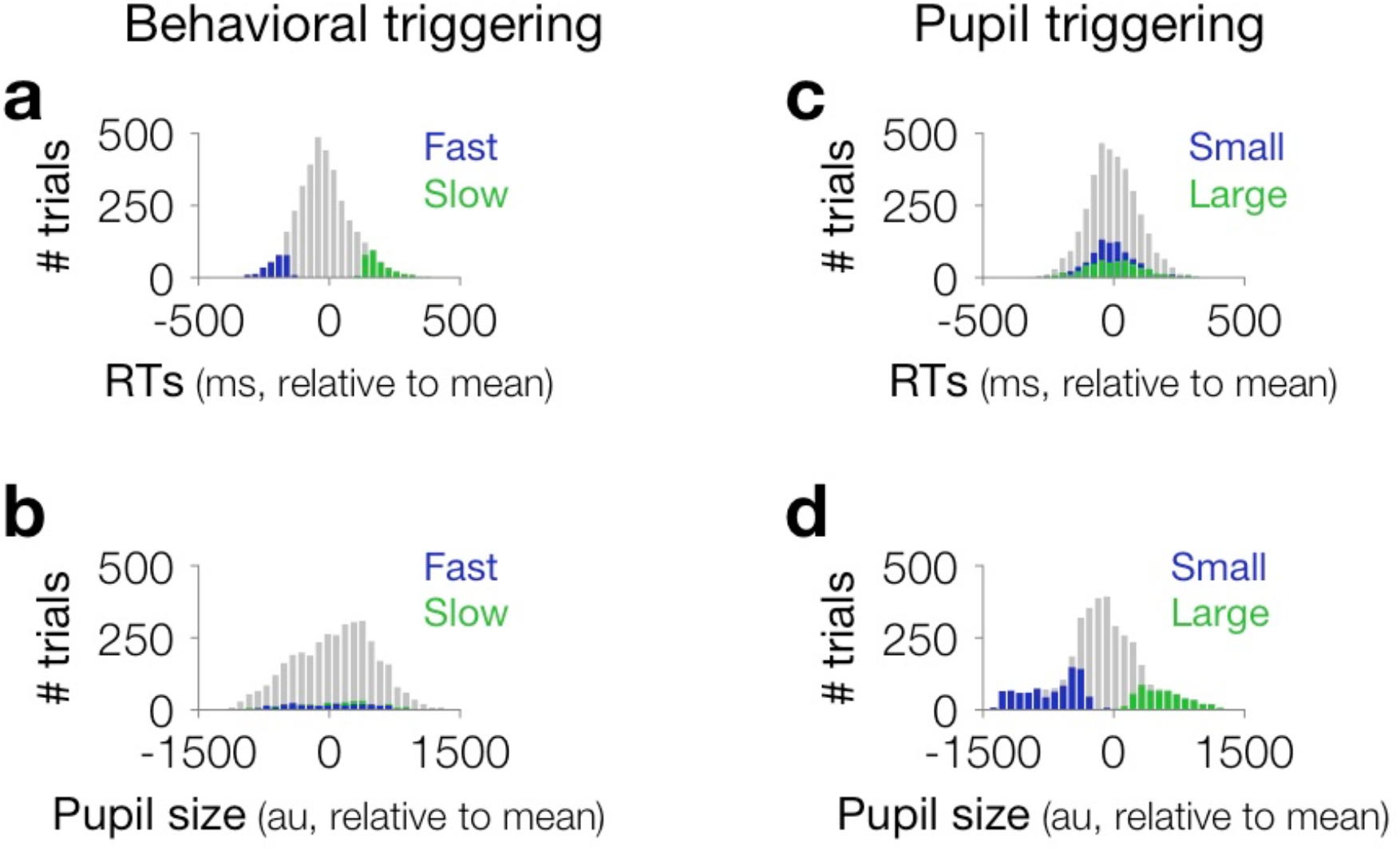
**a** Behavioral triggering tracks extreme differences in RT. A histogram of RTs from all trials of a representative participant in Experiment 1 is depicted in gray. Trials that are eligible for triggering are overlaid, either fast trials (blue) or slow trials (green). **b** Behavioral triggering elicits modest differences in pupil size. A histogram of pupil size from all trials from the same representative participant in Experiment 1 is depicted in gray. Trials that are eligible for triggering are overlaid, either fast trials (blue) or slow trials (green). **c** Pupil triggering elicits modest differences in RT. A histogram of RTs from all trials of a representative participant in Experiment 2 is depicted in gray. Trials that are eligible for triggering are overlaid, either small trials (blue) or large trials (green). **d** Pupil triggering tracks extreme differences in pupil size. A histogram of pupil size from all trials of a representative participant in Experiment 2 is depicted in gray. Trials that are eligible for triggering are overlaid, either small trials (blue) or large trials (green).

#### Pupil triggering

A key advance in Experiment 2 is the development of real-time pupil triggering for this interleaved sustained attention and working memory task. Currently, most attempts to track sustained attentional states in real time have required overt and repetitive behavioral responses. Developing techniques to track attention using pupil size could examine cognitive dynamics covertly and continuously, even without any behavioral response demands (Mathôt et al., 2016). The development of abilities to covertly track sustained attentional state could be especially beneficial in educational or occupational scenarios where even subtle fluctuations of attention could be catastrophic. For pupil real-time triggering, we designed our task based on the pupil signatures observed in Experiment 1. However, other work has implicated that intermediate pupil sizes might be more optimal than high or low (Murphy et al., 2011). Future studies could further improve on this initial demonstration of pupil real-time triggering and can attempt to build upon these findings to develop procedures to prospectively target working memory states.

In this study, we present positive steps forward in exploring the cognitive states and biomarkers underlying lapsing attention. Our findings reveal that lapsing sustained attention states may co-occur with differences in pupil size. This provides additional, complementary insight into the complex physiological signatures that underly lapses of sustained attention (deBettencourt et al., 2015; Esterman & Rothlein, 2019; Rosenberg et al., 2016). In sum, examining pupil size sheds light on moment-by-moment and trial-by-trial fluctuations of sustained attention.

## Author contributions

All authors conceived of the study and contributed to the study design. P.A. Keene and M.T. deBettencourt collected and analyzed the data. P.A. Keene and M.T. deBettencourt wrote an initial draft of the manuscript, which all authors read and edited.

## Acknowledgements

This research was supported by National Institute of Health grant R01MH087214, the Office of Naval Research grant N00014-12-1-0972, and F32MH115597 (M.T.dB.).

